# Differential effects of amplitude-modulated transcranial focused ultrasound on excitatory and inhibitory neurons

**DOI:** 10.1101/2020.11.26.400580

**Authors:** Duc T. Nguyen, Destiny Berisha, Elisa Konofagou, Jacek P. Dmochowski

## Abstract

Although stimulation with ultrasound has been shown to modulate brain activity at multiple scales, it remains unclear whether transcranial focused ultrasound stimulation (tFUS) exerts its influence on specific cell types. Here we propose a novel form of tFUS where a continuous waveform is amplitude modulated (AM) at a slow rate (i.e., 40 Hz) targeting the temporal range of electrophysiological activity: AM-tFUS. We stimulated the rat hippocampus while recording multi-unit activity (MUA) followed by classification of spike waveforms into putative excitatory pyramidal cells and inhibitory interneurons. At low acoustic intensity, AM-tFUS selectively reduced firing rates of inhibitory interneurons. On the other hand, higher intensity AM-tFUS increased firing of putative excitatory neurons with no effect on inhibitory firing. Interestingly, firing rate was unchanged during AM-tFUS at intermediate intensity. Consistent with the observed changes in firing rate, power in the theta band (3-10 Hz) of the local field potential (LFP) decreased at low-intensity, was unchanged at intermediate intensity, and increased at higher intensity. Temperature increases at the AM-tFUS target were limited to 0.2°C. Our findings indicate that inhibitory interneurons exhibit greater sensitivity to ultrasound, and that cell-type specific neuromodulation may be achieved by calibrating the intensity of AM-tFUS.

## Introduction

Owing to the physics of propagating acoustic waves, transcranial focused ultrasound stimulation (tFUS) offers a tantalizing approach to stimulating the brain. The technology to non-invasively stimulate any region of the human brain with millimeter spatial resolution is already in place (1) and has been successfully employed in neurosurgery (2) and to facilitate the targeted delivery of genes and drugs to the brain (3) in the form of high-intensity focused ultrasound (HIFU). When delivered at relatively low intensity, stimulation with ultrasound has been demonstrated to modulate neural activity (4; 5; 6; 7; 8; 9), as manifest by changes in spiking (10; 11), local field potentials (12; 13; 14; 15; 16), fMRI (17; 18; 19; 20; 21; 22; 23; 24), evoked potentials (18; 19; 24; 25; 26; 27; 28; 29; 30; 31), and motor behaviors (10; 15; 16; 24; 30; 32; 33; 34; 35; 36; 37; 38).

Despite abundant evidence for the capability of ultrasound to modulate activity in the central nervous system, there is considerable uncertainty about the mechanism of action (7; 9), and importantly, how to tailor tFUS to produce a desired electrophysiological change (e.g. excite or inhibit activity in a targeted brain region). Thus far, tFUS has been mostly probed as a transient perturbation of either somatosensory or motor activity, where sonication consists of very brief (< 1 s) stimuli of either continuous or pulsed sinusoidal waveform (33). Notable exceptions include recent investigations with longer sonication periods (23) and shorter pulse repetition frequencies (39). Using the conventional paradigm, several studies have reported inhibition of sensory evoked potentials by tFUS (25; 27; 31; 40). Extracellular field potentials are believed to be generated by a superposition of excitatory post synaptic potentials (EPSP) (41), with the amplitude of the evoked potential depending not only on the number and strength of active synaptic currents but also on their geometry and temporal coherence (42). A reduction in evoked potential amplitude can thus reflect either reduced synaptic activity *or* reduced coherence. Moreover, due to the brief nature of inhibitory post synaptic potentials (IPSP), inhibitory interneurons are not believed to significantly contribute to sensory evoked potentials (41). Thus, evoked potential readouts afford only limited insight into the effects of low-intensity ultrasonic stimulation on the activity of different cell types, in particular the relative sensitivity of excitatory and inhibitory neurons to tFUS.

A balance of excitatory and inhibitory signaling is a fundamental property of neural circuits, with imbalances in the excitation-to-inhibition ratio implicated in several psychiatric and developmental disorders (43; 44; 45). Due to their distinct waveforms, it is possible to distinguish the spikes of excitatory and inhibitory cells from recordings of the local extracellular field (46). Several features of the spike waveform such as the trough-to-peak duration or the spike width have been successfully used to separate these two fundamental cell types and to study their response to phenomena such as attentional modulation (47). It is important to understand the influence of tFUS on these two cell classes: first, cell-class specific insight will inform our understanding of the mechanism underlying ultrasonic neuromodulation. Secondly, it is possible that different tFUS waveforms will exert a preferential effect on specific cell types. Indeed, this has been predicted by biophysical models of tFUS that focus on changing membrane capacitance (48; 49). Key parameters expected to have an influence on neuromodulation outcome are the acoustic intensity, pulse repetition frequency, and duty cycle. To date, however, the effect of tFUS parameters on specific classes of neurons has not been empirically probed.

Here we aimed to characterize the effect of ultrasonic stimulation on the two primary cell classes in the hippocampus: excitatory pyramidal cells and inhibitory interneurons. Previous tFUS investigations have employed ultrasonic waveforms where peak intensity is achieved approximately every millisecond (e.g. pulse repetition frequencies in the order of 1 kHz). There is evidence from other forms of neuromodulation, namely deep brain stimulation (DBS) and transcranial magnetic stimulation (TMS), that employing much slower stimulation frequencies allows one to achieve targeted effects (50; 51). We therefore developed a new form of tFUS where a low-frequency rhythm targeting the range of hippocampal activity (40 Hz) was embedded into a high frequency carrier (2 MHz) via amplitude-modulation. Throughout, we refer to this new variant of ultrasonic neuromodulation as “amplitude-modulated tFUS” (AM-tFUS). We measured the local field potential (LFP) and multi-unit activity (MUA) from the rat hippocampus concurrently to the application of AM-tFUS at three intensity levels. To gain insight into the potential role of temperature changes, we subsequently measured the temperature changes induced by AM-tFUS at the same intensities. In what follows, we describe our finding of a cell-class specificity in AM-tFUS, where inhibitory interneurons exhibited sensitivity at low intensities while excitatory pyramidal cells responded to higher intensities. Moreover, these cell-class specific changes were achieved in the absence of significant heating.

## Results

To identify the effect of low-intensity ultrasound on excitatory and inhibitory neurons, we recorded multi-unit activity (MUA) and local field potentials (LFP) from the rat hippocampus before, during, and after three-minutes of AM-tFUS (Fig 1). We employed a within-subjects design, where animals received first sham and then active stimulation at increasing intensities: 1.6, 3.2, and 6.4 W/cm^2^ *I*_spta_ (Fig 1C). In what follows, we refer to these as “low”, ‘‘intermediate”, and “high” intensity, although all three intensities are well below those employed in HIFU, whose effects are thermal in nature. Our primary outcome measures were the changes from baseline in firing rate and LFP power. To determine the magnitude of temperature increase during ultrasonic stimulation, we then conducted follow-up experiments to measure brain temperature in the sonicated region during AM-tFUS.

**Figure 1:**
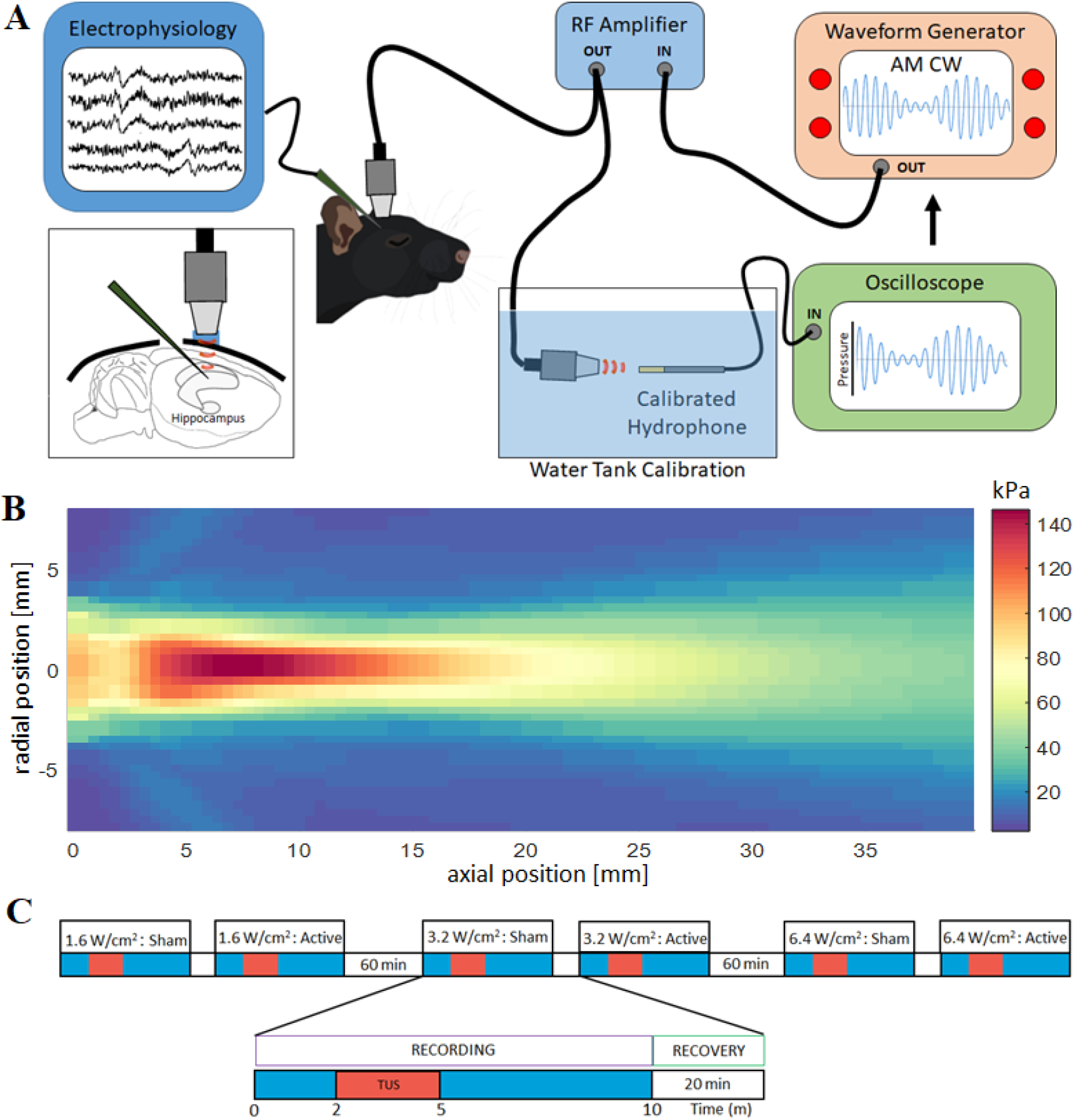
Capturing the effect of AM-tFUS on electrophysiological activity in the hippocampus. (**A**) tFUS was delivered with an ultrasonic transducer coupled to the skull and driven by a function generator and RF amplifier. A multichannel electrode array was advanced into the hippocampus via a craniotomy posterior of the stimulation site. Acoustic intensities were calibrated with a calibrated hydrophone in a water tank. We employed a continuous amplitude modulated (AM) waveform consisting of a 2 MHz carrier and a 40 Hz AM frequency. (**B**) The acoustic beampattern of the transducer employed in this study shows a full-width-half max (FWHM) of 2.4 mm laterally and 10 mm in depth. (**C**) Animals received tFUS at three intensities (“low”: 1.6 W/cm^2^, “intermediate”: 3.2 W/cm^2^, and “high” 6.4 W/cm^2^). To control for spontaneous effects due to passage of time, we performed sham stimulation prior to each active AM-tFUS recording.

### Discriminating excitatory and inhibitory spikes

In order to assess the effect of AM-tFUS on excitatory and inhibitory neurons, we exploited the fact that inhibitory spikes are shorter in duration and produce narrower waveforms (Fig 2A-B). For each identified unit, we measured the trough-to-peak duration and found a bimodal distribution characteristic of a mixture of excitatory and inhibitory spikes (Fig 2C; *p* < 0.001, Hartigan dip test for unimodality). Based on the observed distribution, we selected 900 *μs* as the threshold between inhibitory and excitatory spikes. In our data that was bandpass filtered from 150 Hz to 15 kHz, excitatory spikes had a mean duration of 1165.5±5.2 *μ*s (mean ± sem), while inhibitory spikes had a mean duration of 630.7±10.0 *μ*s. Pooling across the six experimental conditions, we found a total of *n* = 855 excitatory units and *n* = 164 inhibitory units. Visual inspection of the spike waveforms confirmed the presence of two distinct categories of spikes, with putative inhibitory interneurons showing narrower spikes (Fig 2A-B). Importantly, we found good correspondence between our measured firing rates (at baseline) of the two categories of neurons and those reported in prior literature: inhibitory interneurons fired at 15.54±0.38 Hz, while the firing rate of putative excitatory neurons was 1.81±0.06 Hz (means ± sem; Fig 2D). Also recording from the rat hippocampus, Csicsivari et al reported firing rates of 14.1 ± 1.4 Hz in inhibitory interneurons and 1.4 ± 0.01 Hz in excitatory pyramidal neurons (52).

**Figure 2:**
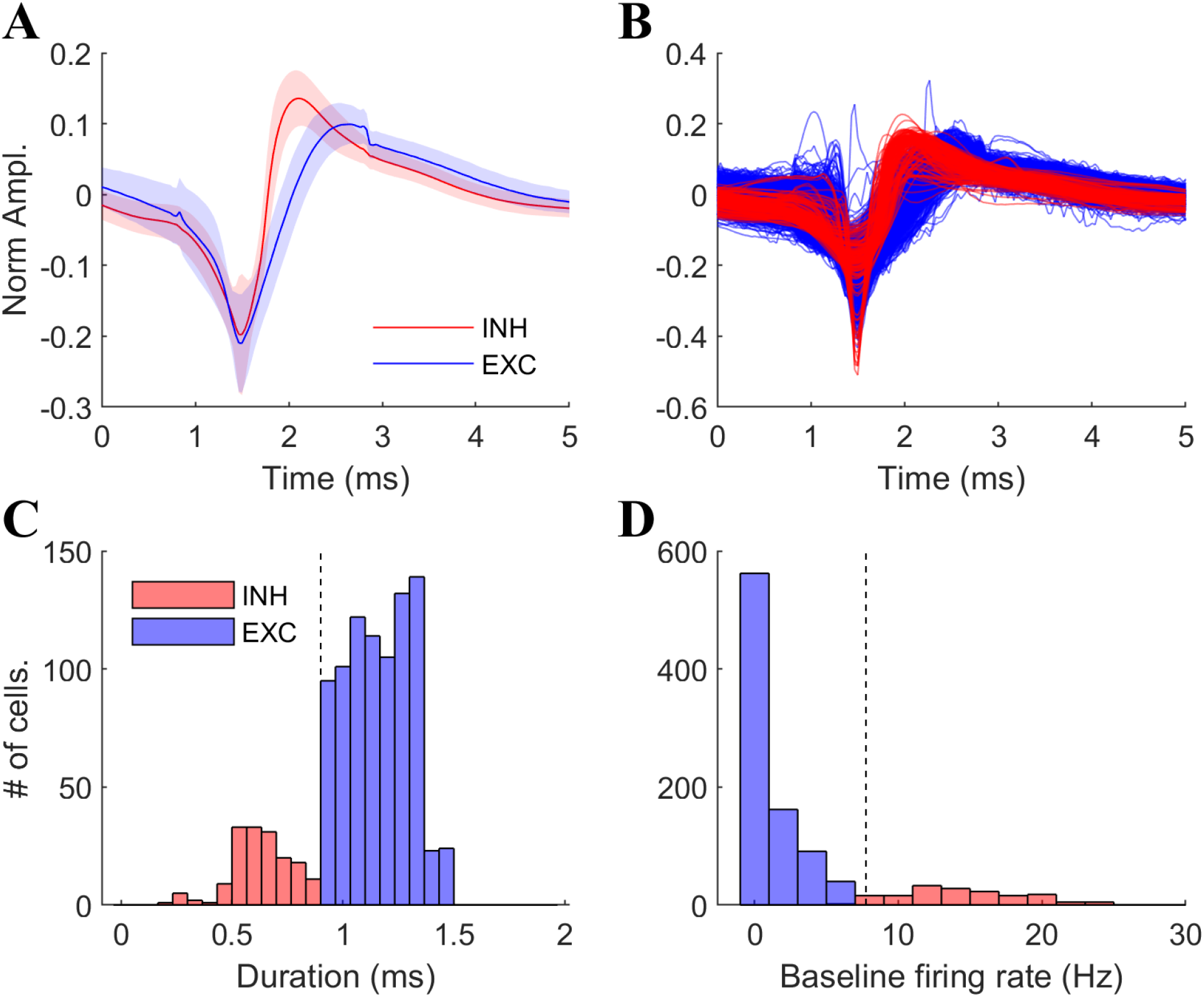
Classification of extracellular spike waveforms into excitatory and inhibitory units. (**A**) Grand-averaged spike waveforms for inhibitory (red) and excitatory (blue) cells. Inhibitory units exhibit waveforms with a shorter trough-to-peak duration (shading indicates sem). (**B**) Mean spike waveforms of individual inhibitory (red) and excitatory (blue) units. (**C**) The distribution of trough-to-peak durations exhibits a bimodal distribution (*p* < 0.001, Hartigan dip test), consistent with a mixture of inhibitory and excitatory spikes. A threshold of 900 *μ*s was employed here to classify cell-type. (**D**) The distribution of baseline firing rates for excitatory and inhibitory units: putative inhibitory interneurons exhibited higher firing rates (15.54 ± 0.34, means ± sem) than that of excitatory pyramidal neurons (1.81 ± 0.06, means ± sem).

### Reduced inhibitory firing at low intensity

During low intensity AM-tFUS, we found a modest but significant reduction in the firing rate of putative inhibitory interneurons relative to sham (Mann Whitney U-test, *z* = −2.11, *p* = 0.035; Fig 3A). The decreased inhibitory firing rate continued in the five minutes following stimulation (*z* = −2.47, *p* = 0.014; Fig 3B). There were no changes in the firing rates of putative excitatory cells at low-intensity, either during or after stimulation (*p* > 0.42; Fig 3C,D).

**Figure 3:**
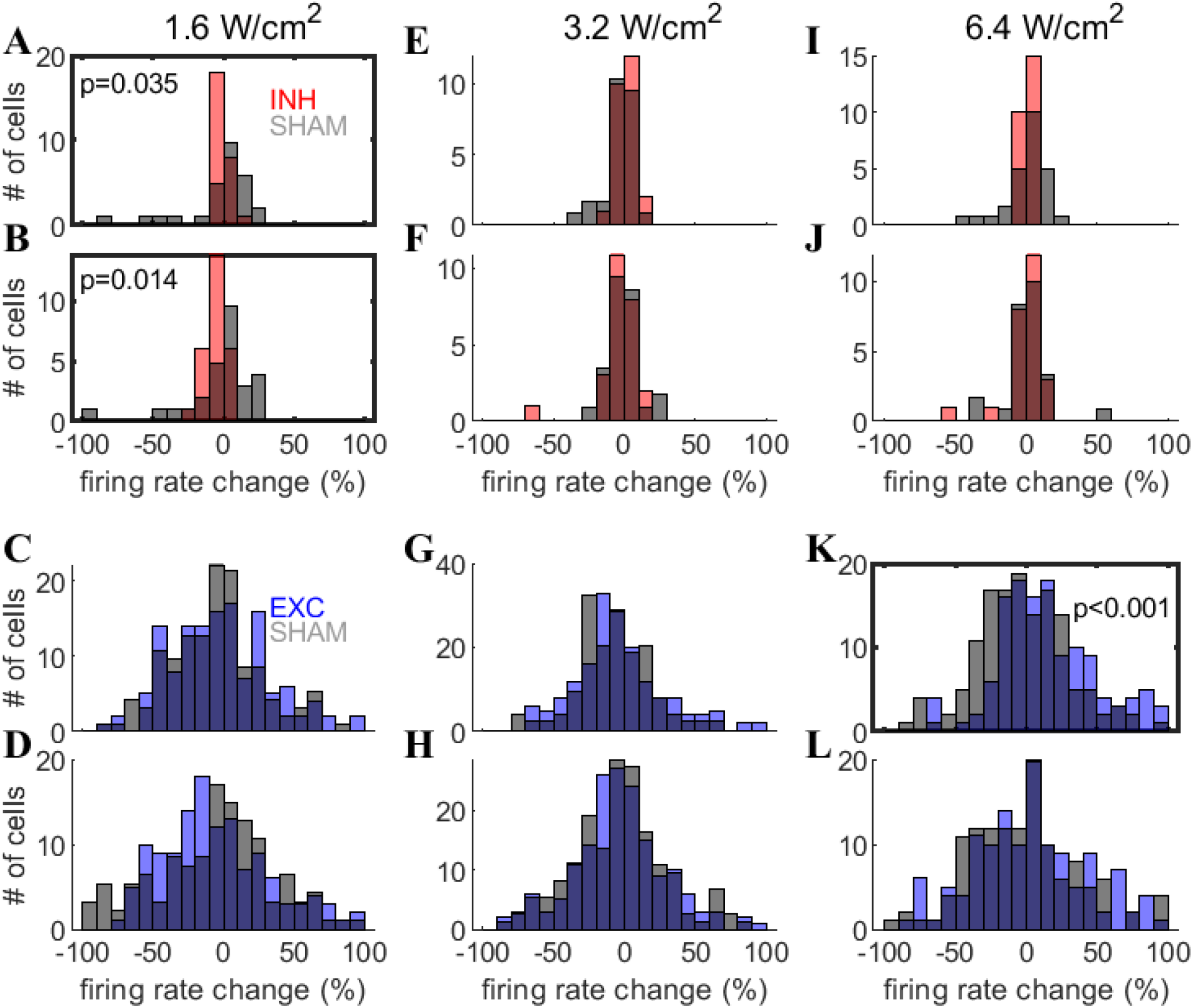
Cell-type specific modulation of firing rate with AM-tFUS. (**A**) At low-intensity, tFUS significantly reduced the firing rate of putative inhibitory interneurons compared to sham (*z* = −2.11, *p* = 0.035, Mann-Whitney U test). (**B**) The reduced inhibitory firing rate continued in the five minute period following stimulation (*z* = −2.47, *p* = 0.014). Firing rate of putative excitatory neurons was unchanged (**C**) during and (**D**) after low-intensity AM-tFUS. (**E-H**) At intermediate intensity, we did not find significant changes in firing rates of inhibitory or excitatory units, either during or after stimulation (all *p* > 0.063). Inhibitory interneurons did not show altered firing rates (**I**) during or (**J**) after high-intensity tFUS (*p* > 0.27). (**K**) On the other hand, we found a large acute increase in the firing rate of putative excitatory units (*z* = 4.87, *p* = 1.1 × 10^−6^) that subsided in the five minutes (**L**) after stimulation (*p* = 0.56).

### No effects at intermediate intensity

We did not resolve an effect on firing rate at the intermediate acoustic intensity for excitatory or inhibitory neurons, either during or after stimulation (Fig 3E-H). Putative inhibitory interneurons exhibited a trend to increase firing during stimulation, but this fell short of reaching significance (*z* = 1.85, *p* = 0.063; Fig 3E).

### Increased excitatory firing at high intensity

During high-intensity stimulation, we found a large increase in the firing rate of putative excitatory neurons (*z* = 4.87, *p* = 1.1 × 10^−6^; Fig 3K). The increased excitatory firing was not sustained in the five minutes following stimulation (*z* = 0.58, *p* = 0.56; Fig 3L). Inhibitory interneurons did not show altered firing rates during or after high-intensity stimulation (*p* > 0.27; Fig 3I-J).

### Stable baseline firing rates across intensities

In order to rule out the possibility of AM-tFUS effects carrying over from one intensity to the next, we measured the baseline firing rate (i.e., the firing rate in the two minutes immediately preceding AM-tFUS onset) and tested for significant differences between intensities. For both inhibitory and excitatory neurons, there were no significant differences in baseline firing rate between low (inhibitory: 17.37 ± 1.28 Hz, excitatory: 1.59 ± 0.16 Hz), intermediate (inhibitory: 16.89 ± 1.16 Hz, excitatory: 1.79 ± 0.16 Hz), and high-intensity (inhibitory: 15.52 ± 0.79 Hz, excitatory: 1.90 ± 0.17 Hz). All pairwise comparisons failed to detect a significant effect (inhibitory, *p* > 0.37; excitatory: *p* > 0.11; Mann Whitney U-test).

### Bidirectional modulation of theta power

To probe AM-tFUS driven changes in local synaptic activity, we computed the power spectrum of the LFP during the baseline, stimulation, and poststimulation segments (Fig 4A-B; depicted as averages across *n* = 20 animals). Spectra exhibited characteristic 1/f shapes with inflections near the theta (3-10 Hz) region. To identify significant changes in power spectrum, we computed the percentage of spectral power change from baseline at each frequency during (Fig 4C-E) and after (Fig 4F-H) stimulation. This revealed a bidirectional effect, with low-intensity AM-tFUS producing an acute *decrease* of theta power relative to sham (Fig 4C; significant cluster from 3.2 to 9.2 Hz, 23% reduction relative to sham; *p* = 0.004, paired two-tailed Wilcoxon sign rank test, *n* = 20, cluster corrected with a permutation test randomizing active and sham samples; curves show mean ± sem across animals), while high-intensity AM-tFUS produced a relative *increase* of theta power during stimulation (Fig 4E; 43% increase at a cluster from 4.7-8.4 Hz; *p* = 0.004). There were no significant acute changes found at intermediate intensity (Fig 4D). The results here are shown for contacts in the dentate gyrus region. Similar results were found for areas CA1 and CA3 (Fig S1 and S2).

**Figure 4:**
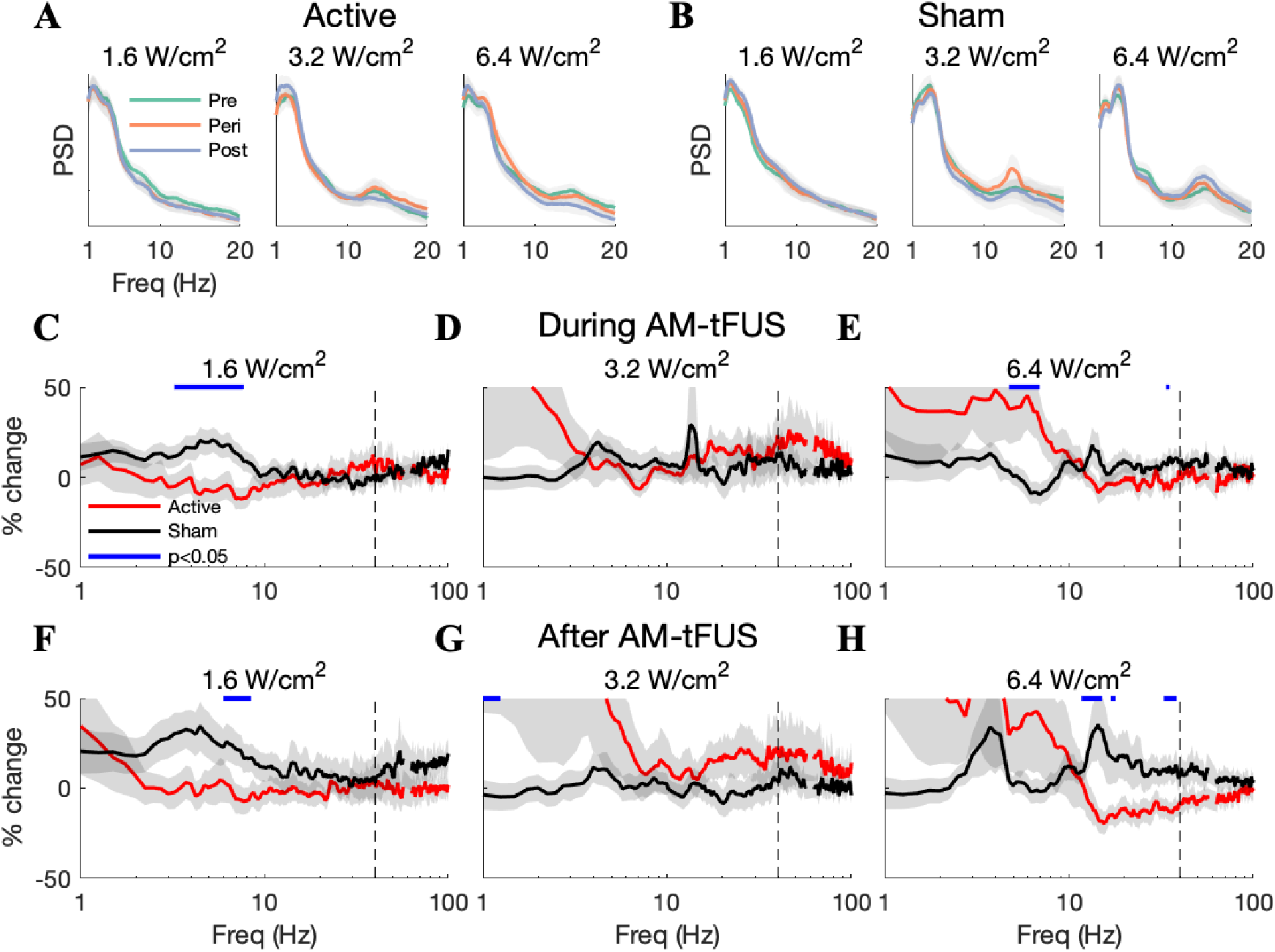
AM-tFUS modulates LFP spectral power in an intensity and frequency-dependent manner. (**A**) Power spectral densities before (green), during (red), and after AM-tFUS at low (left), intermediate (middle), and high (right) intensity. (**B**) Same as (A) but now shown for the sham stimulation recordings. The characteristic 1/f LFP spectrum, in addition to an inflection over the theta (3-10 Hz) band was observed for all conditions. (**C**) During low intensity AM-tFUS, a significant reduction in spectral power was observed at a cluster in the theta band (cluster indicated with blue line, 23% reduction, *p* = 0.004, Wilcoxon sign rank test, *n* = 20, cluster corrected with a permutation test employing scrambling of active and sham assignment). (**D**) No significant clusters were observed during AM-tFUS at intermediate intensity. (**E**) At high-intensity, a significant increase in power relative to sham was observed at a cluster in the theta band (43% increase, *p* = 0.004). (**F**) The reduced theta power at low-intensity AM-tFUS was sustained after stimulation (*p* = 0.022). (**G**) A significant increase in the delta (1-3 Hz) band was observed following stimulation at intermediate intensity (*p* = 0.012). (**H**) Although the increase in theta power was not sustained following high-intensity AM-tFUS, we found significant power reductions in several higher frequency clusters in the 12-50 Hz region after high-intensity AM-tFUS (*p* < 0.048). Shaded error bars denote sem.

The reduction in theta power at low-intensity was sustained during the five minutes following stimulation (Fig 4F; significant cluster from 6.0 Hz to 9.9 Hz, *p* = 0.022). On the other hand, the acute theta increase at high-intensity did not persist in the period after stimulation (Fig 4H; no significant cluster in the theta range). Interestingly, we also found effects that *only* manifested in the post-stimulation period. At intermediate intensity, a significant post-stimulation increase was found between 1 and 3 Hz (Fig 4G, *p* = 0.012), while significant decreases at high-intensity were found in the following clusters (Fig 4H): 11.7-16.6 Hz (*p* = 0.016), 16.9-19.4 Hz (*p* = 0.018), 32.8-40.0 Hz (*p* = 0.0040), 47.2-50.4 Hz (*p* = 0.048).

To further probe the bidirectional effect of AM-tFUS on activity in the theta band, we computed the mean spectral power in the entire 3-10 Hz range before, during, and after stimulation, and then measured the change from baseline for all animals (Fig 5A-C; shown for the dentate gyrus). We conducted a two-way repeated-measures (*n* = 20) ANOVA with intensity and condition (active/sham) as factors and the percent change from baseline as the dependent variable. We found a significant interaction between intensity and condition during but not after AM-tFUS (during: *F*(2) = 5.76, *p* = 0.004; after: *F*(2) = 1.70, *p* = 0.19), confirming that AM-tFUS modulated the dynamics of theta power in an intensity-dependent manner, at least during stimulation. Follow-up pairwise tests identified a significant theta reduction both during and after low-intensity AM-tFUS (Fig 5A, during AM-tFUS: −19.5 % ± 7.66%, *p* = 0.017, *n =* 20, Wilcoxon sign rank test; after AM-tFUS: −24.1 % ± 11.9%, *p* = 0.023; means ± sem). At high-intensity, a significant increase in theta power was identified during AM-tFUS (Fig 5C, 29.6 % ± 13.0%, *p* = 0.048, *n* = 20). There was no significant change in theta power at intermediate intensity, either during or after stimulation. Similar changes in theta power were observed in the CA1 and CA3 subregions (Figs S3 and S4).

**Figure 5:**
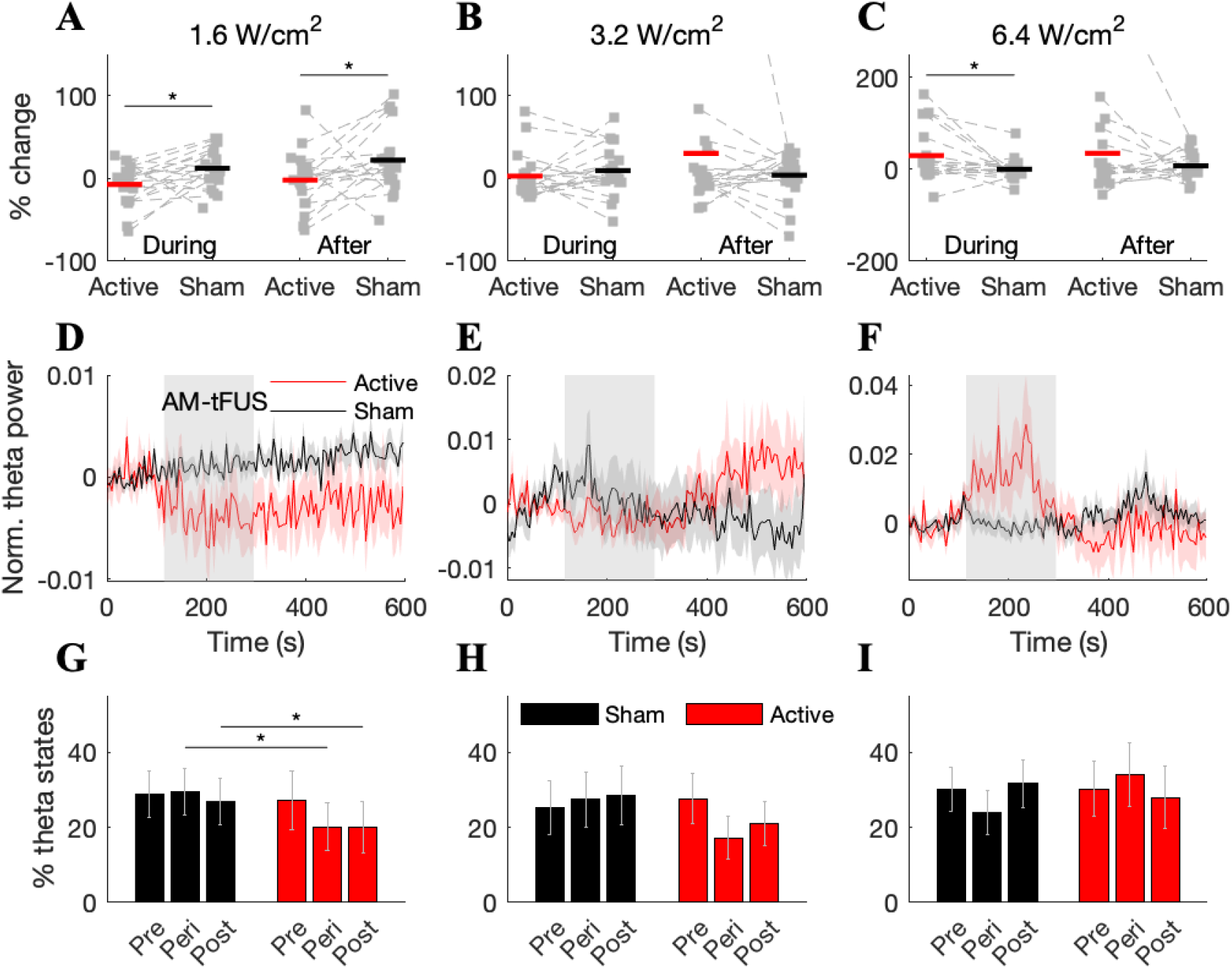
AM-tFUS produces a bidirectional modulation of theta power. To probe the effect on the theta band, we measured the change from baseline in the 3-10 Hz band for both active and sham stimulation. (**A**) At low-intensity, AM-tFUS significantly reduced theta power during and after stimulation, relative to sham (during: −19.5 ± 7.7%, *p* = 0.017; after: −24.1 ± 11.9%, *p* = 0.023, *n* = 20, Wilcoxon sign rank test). (**B**) Theta power was unchanged during or after intermediate intensity AM-tFUS. (**C**) High-intensity AM-tFUS produced a significant increase of theta power during (29.6 ± 13.0%, *p* = 0.048) but not after stimulation. (**D**) The time course of theta power, measured in five second windows, exhibited a sharp decrease near low-intensity stimulation onset. (**E**) Same as (D) but for intermediate intensity. (**F**) High-intensity AM-tFUS evoked an immediate increase in theta power at stimulation onset that subsided following stimulation. (**G**) We measured the proportion of time spent in so-called “theta” states, marked by increased levels of the ratio of theta-to-delta power. We observed a significant decrease in the proportion of time spent in theta during and after low-intensity AM-tFUS (during: *p* = 0.018, after: *p* = 0.010, Wilcoxon sign rank test, *n* = 20). However, no change in the proportion of time spent in theta was observed (**H**) during or (**I**) after stimulation at intermediate or high intensity.

To gain insight into the dynamics of the observed theta-band changes, we measured theta power in a time-resolved fashion (i.e., 5 second windows). At low intensity, the sham stimulation time course showed a steady increase throughout the 10-minute recording period, while active AM-tFUS produced an immediate sharp reduction near stimulation onset that was sustained after stimulation (Fig 5D). On the other hand, high-intensity AM-tFUS produced an immediate increase at stimulation onset that was abolished following stimulation (Fig 5F).

In the awake state, the rat hippocampus is known to alternate between “theta” and “non-theta” states, which loosely correspond to active (locomotion and REM sleep) and idle states (53; 54). Analogues of these states have been observed under anesthesia (55). We suspected that AM-tFUS may modulate the proportion of time spent in theta. Relative to sham, we found a significant reduction in theta states at low-intensity (Fig 5G; during: *p* = 0.018; after: *p* = 0.010; *n* = 20; paired two-tailed Wilcoxon sign rank test; shown here for the dentate gyrus). No significant differences in proportion of time spent in theta were found at the intermediate or high-intensities (Fig 5H-I).

### Temperature increases are limited to 0.2°C

Given the three-minute sonications employed in this study, it is important to determine whether heating of the brain occurred, and if so, to assess its contribution to the observed neural changes. We therefore conducted follow-up experiments (*n* = 8) that replicated the main experiments, except that the recording electrode was replaced with a thermocouple (Fig 6A). As expected, temperature increased monotonically during stimulation, with the largest increases observed during high-intensity AM-tFUS (Fig 6B-D). However, the magnitude of the increases was relatively small for all intensities: at the end of the three-minute stimulation period, the temperature change from baseline was Δ*T* = 0.05 ± 0.14, 0.12 ± 0.19, and 0.20 ± 0.18°C, for low, intermediate, and high-intensity stimulation, respectively (mean ± sem across *n* = 8 animals). The highest instantaneous temperature increases during AM-tFUS (relative to baseline) recorded were 0.33, 0.59, and 0.67°C for low, intermediate, and high-intensity, respectively.

**Figure 6:**
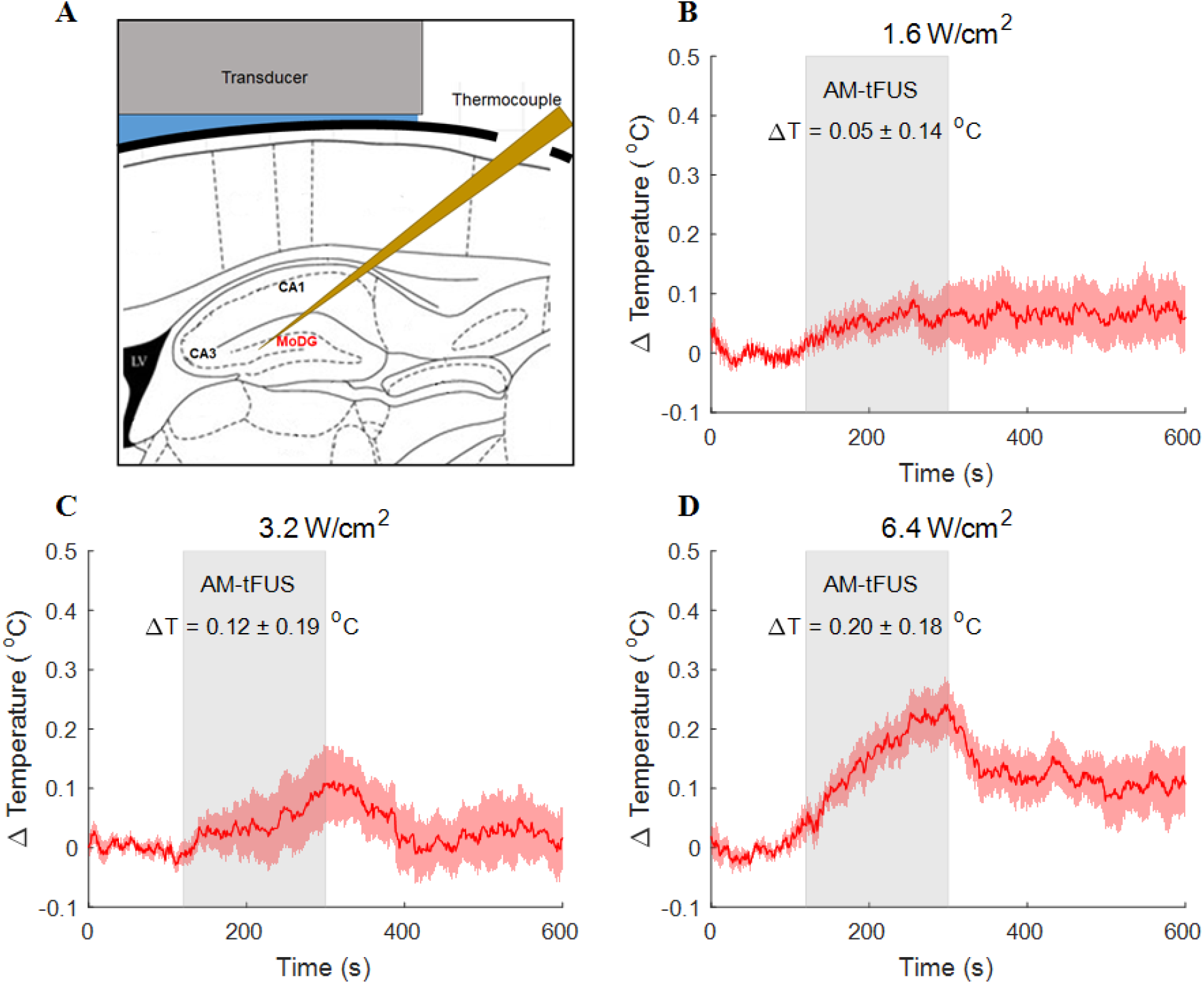
Temperature increases at the sonicated region are limited to 0.2°C. (**A**) To determine the temperature increase produced by AM-tFUS, we stimulated the hippocampus but now with a thermocouple inserted into the focus of the ultrasonic beam. Temperature was recorded before, during, and after AM-tFUS. As expected, temperature increased monotonically during stimulation. However, the observed temperature increases were low, with mean changes at the end of stimulation of (**B**) 0.05 ± 0.14°C, (**C**) 0.12 ± 0.19°C, and (**D**) 0.20 ± 0.18°C at low, intermediate, and high intensity, respectively (means ± sem, *n* = 8). Shaded error bars denote sem.

## Discussion

Here we proposed and tested a new form of ultrasonic neuromodulation where a low-frequency rhythm is embedded into a high-frequency continuous waveform via AM. By recording and classifying extracellular spikes during AM-tFUS, we found that putative inhibitory interneurons reduced their firing during low-intensity stimulation while excitatory neurons responded at higher intensity by increasing firing rate. No effects were observed at an intermediate intensity. Congruent with the firing rate results, the power of theta band oscilations decreased at low intensity while increasing at high intensity. The temperature of the sonicated region rose by an average of 0.2°C at high-intensity. These findings provide the first experimental evidence for the notion that specific cell types can be targeted by appropriate selection of AM-tFUS parameters.

We found congruence between the measured changes in neuronal firing rates and those in the LFP theta power. Inhibitory interneurons have been previously reported to have a causal role in the generation of theta rhythms, as mice bred with impaired inhibitory synaptic activity exhibit lower theta, but not gamma, power (56). Moreover, inhibitory interneurons exhibit spike timing that is phase locked with theta cycles (57; 58; 59; 60). Therefore, one interpretation of the findings at low-intensity stimulation is that AM-tFUS impaired action potential generation in inhibitory interneurons, consequently producing less field activity (or perhaps less coherence) in the theta range. In this scenario, inhibitory interneurons exhibit reduced firing and the power of theta oscillations would be decreased, consistent with what was observed here. Secondly, theta oscillations in the rat hippocampus are generated from the rhythmic excitatory input from the entorhinal cortex to the apical dendrites of pyramidal cells in the CA1 and CA3 regions of the hippocampus (61; 62; 63). Exogenous stimulation of these pyramidal cells with AM-tFUS would be expected to then increase the power of theta oscillations, consistent with what was observed here at high-intensity AM-tFUS. Taken together, the results of the firing rate analysis are aligned with the measured changes in theta rhythms.

Interestingly, we did not find significant changes in firing rate or LFP power at the intermediate intensity. This suggests that the observed modulations of inhibitory interneurons at low intensity and excitatory pyramidal cells at high-intensity reflect two separate phenomena, with potentially distinct biophysical substrates. The proposed mechanisms of neuromodulation by tFUS can be partitioned into mechanically and thermally mediated changes (7; 9). The mechanical pathways include changes in the ion channel permeability (7; 64; 65; 66) that occur when the acoustic radiation force causes a mechanical strain on the plasma membrane, and indirect changes to membrane potential due to changing membrane thickness or capacitance (48; 49). The thermal pathway involves capacitive currents induced by temperature rises that promote neural excitation (67; 68; 69) or changes in the permeability of heat-sensitive ion channels (70; 71). Our finding of opposing effects at low and high intensity with no effect at intermediate intensity may be explained by a primarily mechanical effect at the low intensity and a primarily thermal effect at high intensity. This view is supported by the very limited temperature increase of 0.05° C at low-intensity AM-tFUS – given the low magnitude of this temperature gradient, the reduction in inhibition and reduced theta power that we found indicate a mechanical effect. Consistent with the view of a thermal effect producing increased excitation, the excitatory effect at high intensity did not remain after stimulation was turned off (i.e., when the mild heating subsided). It is thus possible that the mild temperature changes observed at high-intensity AM-tFUS altered the permeability of heat-sensitive ion channels and increased the pyramidal cells’ excitability. On the other hand, the mean temperature change observed here at high-intensity (0.2°C) is an order of magnitude lower than that which has been reported to produce inhibition of somatosensory evoked potentials (31). Moreover, this level of temperature elevation is well within the range of physiological fluctuations of brain temperature, including the temperature changes produced by neural activation itself (72; 73; 74). Specifically in the rat hippocampus, temperature increases of 1.5 °C have been measured during active exploration (75), likely reflecting the exothermic nature of brain metabolism. Therefore, further studies are needed to pinpoint the mechanisms explaining the intensity-dependent effects on hippocampal neurons found here.

Given the differential pattern of modulation at low acoustic intensity, our findings suggest that inhibitory interneurons are more sensitive to weak ultrasonic stimuli. This sensitivity may be related to the presence of ion channels expressed by interneurons (76) that are more easily activated with mechanical forces. Inhibitory interneurons also differ from excitatory pyramidal cells in their morphology, and this may mediate some of the cell-type specific responses to AM-tFUS. For example, the large arbors of basket and chandelier cells may be better suited to transducing the mechanical force into changes in electrical properties. Regardless of the mechanism governing this differentiation, it may allow investigators to target specific cell types with appropriately selected acoustic intensity. In practice, this will require knowledge of dose-response characteristics that extend beyond the three intensities tested here.

The recently proposed neuronal intramembrane cavitation excitation (NICE) model (49) has predicted the existence of a cell-type specificity achievable by varying the duty cycle and intensity of conventional tFUS, with qualitative agreement between the model predictions and the findings of previous investigations. The cell-type specificity predicted by NICE is rooted in the lower activation threshold of low-threshold spiking (LTS) inhibitory interneurons to the ultrasonic stimulus, particularly at low duty cycles. The disparity in activation threshold of LTS, fast-spiking interneurons, and pyramidal cells, combined with their connectivity patterns, led to their prediction of an inhibition zone at low intensity, a transition zone at intermediate intensity, and an excitatory zone at high intensity. The intensity dependent, cell-type specific effect predicted by NICE certainly resembles the effects found here *in vivo*. In contrast to the unipolar effects on firing rate reported in NICE (i.e., cells become more excitable during conventional tFUS), here we found empirical evidence that fast-spiking inhibitory interneurons *reduce* their firing rate to low-intensity AM-tFUS. It is possible that the discrepancies between the predictions of NICE and the findings here are a result of a mechanism other than intramembrane cavitation and changing membrane capacitance: for example, acoustic radiation force (77). Another notable difference between the two studies is the use of a sustained AM waveform here, whose fluctuating envelope may lend itself well to periodic displacement of neuronal membranes. Another possible source of variation is our examination of the hippocampus, a region mainly comprised of fast-spiking inhibitory interneurons and pyramidal cells with a connectivity pattern that is generally different than that of the cortical and thalamic structures modeled by NICE.

Our experimental findings underscore the potential of AM-tFUS in producing a diversity of effects on neuronal populations simply by changing the parameters of the ultrasonic stimulus. Coupled with ultrasound’s capability of deep and focused stimulation of the intact human brain, tFUS represents a technique with immense potential as a therapeutic tool and circuit mapping technique. To realize this potential, it will be important to carry out comprehensive parameter sweeps in the awake brain (35), as this will mitigate the presence of spontaneous fluctuations in brain state during anesthesia, and the potential mismatch between tFUS effects on an anesthetized versus awake brain. For example, despite its desirable generation of a long-lasting stable plane of anesthesia, urethane has been previously found to affect multiple neurotransmitter systems (78). Furthermore, in order to disambiguate the influence of mechanical and thermal forces in shaping neuronal changes by tFUS, it will be necessary to conduct experiments that combine electrophysiology with measurements of either mechanical displacement (for example, harmonic motion imaging (79)) or brain temperature (80). Such multimodal approaches can then be combined with tools from statistical inference to provide estimates of the relative contributions of mechanical and thermal forces to the neuronal changes discovered here with AM-tFUS.

## Methods

Data were obtained from 28 (electrophysiology: *n* = 20, thermometry: *n* = 8) adult male Long Evans rats weighing at least 350g (420.54 ± 40.35g, mean ± sd). All experimental procedures were approved by the Institutional Animal Care and Use Committee of the City College of New York, City University of New York.

### Transcranial focused ultrasound

The amplitude modulated waveform (AM frequency 40 Hz, 100% AM depth, carrier frequency 2.0 MHz, sinusoidal) was generated by a function generator (Keysight 33500B Series). The output of the function generator was fed into the input of an RF Amplifier (Electronics and Innovation, 40 W). The amplifier provided the drive voltage into the ultrasonic transducer (Ultran KS25-2 immersion transducer, 2 MHz, 6.25 mm active diameter). The transducer was mounted onto a micromanipulator arm of the stereotaxic frame (David Kopf Instruments) and coupled to the rat skull with ultrasonic coupling gel. To implement sham stimulation, the transducer was translated vertically by 1 cm such that no pressure was delivered to the skull. All other procedures were performed consistently with that of active stimulation.

### Experimental design

A within-subjects design was employed where all animals received three intensities of AM-tFUS, each performed in both active and sham configurations. Similar to the design of Helfrich et al (81), all animals received sham stimulation prior to active stimulation in order to mitigate potential outlasting effects from AM-tFUS leaking into the sham session. The order of the intensities was fixed for all animals: 1.6 W/cm^2^, 3.2 W/cm^2^, 6.4 W/cm^2^. A 20 minute interval was added between the end of the sham recording and the beginning of the active recording. A one hour interval was added between successive intensities in an attempt to achieve consistent baselines (due to time limitations, six rats received only 20 minutes between intensities).

### Acoustic intensity calibration

Ultrasonic pressures were calibrated in a water tank with a calibrated hydrophone (Onda Corporation). Pressures were determined in both free field as well as with a model rat skull placed between the transducer and hydrophone. From this, it was determined that the skull attenuates the acoustic pressure to a value that is 2/3 of the free-field pressure. We then determined input voltages on the function generator such that the spatial-peak temporal-average intensity (*I*_spta_) for a continuous waveform traversing the skull was equal to 2.5 W/cm^2^, 5.0 W/cm^2^ and 10.0W/ cm^2^, for the “low”, “intermediate”, and “high” intensities employed here. After taking into account the AM nature of the waveform, these values are reduced to 1.6, 3.2, and 6.4 W/cm^2^, yielding the final intensity values reported in the main text. In order to generate the acoustic beampattern on a fine spatial grid, we employed the k-Wave toolbox (82) to simulate our transducer’s pressure distribution (Fig 1B).

### Anesthesia and Surgery

Prior to surgical experimentation, selected animals weighing at least 350g were fasted for 12-14 hours to increase urethane absorption. On the day of surgical experimentation, animals were placed in an induction chamber and induced with gaseous isoflurane at 3% (L/min). Animals were removed from the induction chamber and a nose cone was attached so that the dorsal hair could be shaved in preparation for the craniotomy. Animals were then placed on a stereotaxic frame (David Kopf Instruments) with earbars securing the head. The isoflurane concentration was then reduced to 2% (L/min). A cross incision was performed over the dorsal skull to expose the cranium and skull landmarks. The distance from bregma to the interaural line was measured so that anteroposterior (AP) coordinates could be adjusted to account for differences in animal size. In order to position the ultrasonic transducer such that the beam focus coincided with the targeted right hippocampus, we marked the skull at −3.5 AP and +2.5 mediolateral (ML). We targeted a depth of −3.5 mm dorsoventral (DV) from the skull surface. Similarly, the center of the craniotomy was marked at −7.5 AP and +2.5 ML (directly posterior of the transducer). This allowed enough clearance between the edge of the transducer and the border of the craniotomy.

A 2 mm by 2 mm craniotomy was performed over the marked area, followed by the removal of the dura. A small titanium screw was implanted into the skull (left hemisphere) to provide an electrical ground for the electrophysiological recordings. The concentration of isoflurane was further reduced to 1% (L/min) and a urethane cocktail (1.5g/kg diluted with 2.5ml/g saline, divided into 3-4 doses with one dose administered every 10 minutes) was administered via intraperitoneal injection. After the final urethane injection, isoflurane was again lowered to 0.5% (L/min) to allow for urethane absorption and anesthesia transition. After 30 minutes, isoflurane was discontinued and a 90-minute period was allowed for complete expulsion of isoflurane and to achieve a stable anesthesia plane prior to the experiment.

### Electrophysiology

Multi-unit activity (MUA) and local field potentials (LFP) were recorded with a 32-channel silicon electrode array (NeuroNexus A32, 100 *μ*m spacing between adjacent contacts). Signals were recorded with a digital acquisition system (NeuroNexus SmartBox) at a sampling rate of 30 kHz. The probe was placed into the center of the craniotomy at an angle of 53° from vertical (angled towards the posterior), and then advanced 6 mm so that the contacts sampled multiple subregions of the hippocampal formation, including CA1, CA3, and the dentate gyrus. Electrophysiological recording commenced two minutes before the onset of AM-tFUS, continued throughout the three-minute stimulation period as well as an additional five minutes post-stimulation, resulting in 10-minute data records for each experimental condition.

### Thermometry

We performed thermometry in *n* = 8 animals. Sham stimulation was omitted from the thermometry experiments. The procedures for the thermometry experiment mimicked those of the main electrophysiology study. In place of the electrode, a type “T” thermocouple needle (23 AWG, Thermoworks) was inserted into the center of the craniotomy at a 53° angle to the vertical. The thermocouple was advanced 4.5 mm into the brain, such that the tip of the thermocouple coincided with the center contact of the electrode. Temperature was sampled at 50 Hz over a recording duration of 10 minutes. AM-tFUS was applied two minutes after recording onset and had a duration of three minutes. An interval of 20 minutes was included between successive recordings to allow any temperature increases to subside prior to the start of the next recording.

### LFP analysis

All processing was performed offline in the Matlab software package (Mathworks, Release 2019a). Data was bandpass filtered to the 1-250 Hz band with a second-order Butterworth filter and then downsampled to 500 Hz. A series of notch filters were then applied to remove 60 Hz noise and its first four harmonics. To remove gross artifacts from the data, we applied the robust principal components analysis (robust PCA) algorithm, which decomposes a matrix into low-rank and sparse components (83). Due to the smoothness of volume conducted signals, the sparse component is expected to be artifactual and was thus removed from the data. Data was then transformed into the frequency domain by performing the Thomson multi-taper spectral analysis technique (84) with time-bandwidth product of 4. In order to account for varying power levels across animals, all spectra were normalized by dividing by the total spectral power measured in the pre-stimulation period of each animal’s first condition.

Spectra were further downsampled in frequency to 400 samples between 1 and 100 Hz via linear interpolation (we did not analyze spectral energy above 100 Hz). Spectral values for any frequency greater than 57 and less than 63 Hz were marked as missing data due to possible contamination from line noise. Signals were then averaged across contacts belonging to anatomically defined, contiguous regions (dentate gyrus, CA1, and CA3) – in the main text (Figs 4 and 5), we show results for the dentate gyrus. The dependent variable was the change in spectral power from baseline. When depicting time-resolved spectral power (Fig 5D-F), we employed a five-second sliding window with zero overlap among successive windows. Moreover, the time courses were normalized by subtracting the mean power measured in the two-minute pre-stimulation period from every time point.

To perform the analysis of “theta states” in an unbiased manner, we employed the criteria proposed by Shinohara et al (54). A time window was classified as a theta state if the ratio of theta-to-delta power exceeded 0.6, where the theta band was defined as 3.5-7 Hz, and the delta band as 2-3 Hz.

### MUA analysis

In order to perform spike detection and spike sorting, we employed the “Kilosort 2” Matlab toolbox (85). This technique forms a generative model of the extracellular voltage, learns a spatiotemporal template of each spike waveform based on the singular value decomposition, and employs multiple passes through the data to yield well-separated units. We employed the default parameters provided by the developers, as described in the “StandardConfig_MOVEME.m” file provided on https://github.com/MouseLand/Kilosort. The applied highpass filter had a cutoff frequency of 150 Hz. Channels with a firing rate of less than 0.1 Hz were excluded from analysis. Detection of a spike required a negative deflection of at least 6 standard deviations from the channel mean. In a few recordings (i.e., 6), a very low number of spikes was observed. In these cases, the spike detection threshold was lowered to 5 standard deviations, and the procedure was re-run. Due to limitations on the maximum file size, spike sorting was performed independently on the data of each intensity/condition. Thus, six independent runs of the procedure were performed for each animal.

The results of the spike sorting were post-processed to retain only those units that exhibited physiological spiking behavior. For each unit, we measured the basal firing rate and the trough-to-peak duration of the unit’s template waveform. Any unit with a firing rate below 0.1 Hz was excluded from analysis. Units with a trough-to-peak duration shorter than 100 *μ*s or longer than 1500 *μ*s were also excluded from the analysis. To remove units with clearly artifactual waveforms, we excluded units that did not exhibit their trough within ± 250 *μ*s of the threshold crossing, or those that exhibited a *positive* peak within ± 167 *μ*s of the threshold crossing.

In order to classify units into excitatory and inhibitory, we employed *a priori* knowledge of both the spike duration and the basal firing rate. A unit was classified as excitatory if its trough-to-peak duration exceeded 900 *μ*s and if its basal firing rate was less than 7.75 Hz. Conversely, a unit was classified as inhibitory if its trough-to-peak duration was less than 900 *μ*s and its basal firing rate exceeded 7.75 Hz. The firing rate boundary was obtained from the results of Csicsvari et al (52), who found that excitatory pyramidal neurons in the rat hippocampus fire at 1.4 Hz, while inhibitory interneurons fire at 14.1 Hz. The firing rate threshold was thus established as 1.4 + (14.1 – 1.4)/2 = 7.75 Hz. The spike duration threshold was determined by observing the distribution of trough-to-peak duration and noting a “dip” in the bimodal histogram at this value (Fig 2C).

### Thermometry analysis

Raw temperature data were imported into MATLAB via custom scripts. We normalized the temperature time series by subtracting the mean temperature in the two-minute baseline period.

### Statistical testing

Unless otherwise specified, testing for statistical significance was carried out by comparing the dependent variable measured with active stimulation against that observed with sham stimulation. When testing for significant differences in firing rates, we employed the Mann-Whitney U Test (also sometimes termed the Wilcoxon rank sum test). Note that the number of detected units varied with the condition, thus necessitating a non-paired test. When testing for significant differences in LFP power, we employed the Wilcoxon sign rank test, as these measurements were collected on a within-subjects level.

To control for multiple comparisons in the analysis of spectral power changes (Fig 4), we employed a cluster correction technique. A significant frequency cluster was defined as a set of at least 8 consecutive frequency bins (i.e., at least 2 Hz) with more than half of those bins showing a significant effect (uncorrected *p* < 0.05 with the Wilcoxon sign rank test). Significant clusters were detected and then corrected for false positives by employing a permutation test that formed mock spectra where the assignment of active and sham AM-tFUS was scrambled. That is, we formed 500 mock replicates of both active and sham spectra, and then took the difference. Clusters were marked significant after correction if the true difference exceeded 95% of the mock differences.

When testing for significant differences in the proportion of time spent in “theta”, we compared raw proportions without computing a change from baseline. In other words, we compared the proportion of time spent in theta during active AM-tFUS with that measured during sham AM-tFUS, and repeated this for the post-stimulation segment.

---

**Figure S1:**
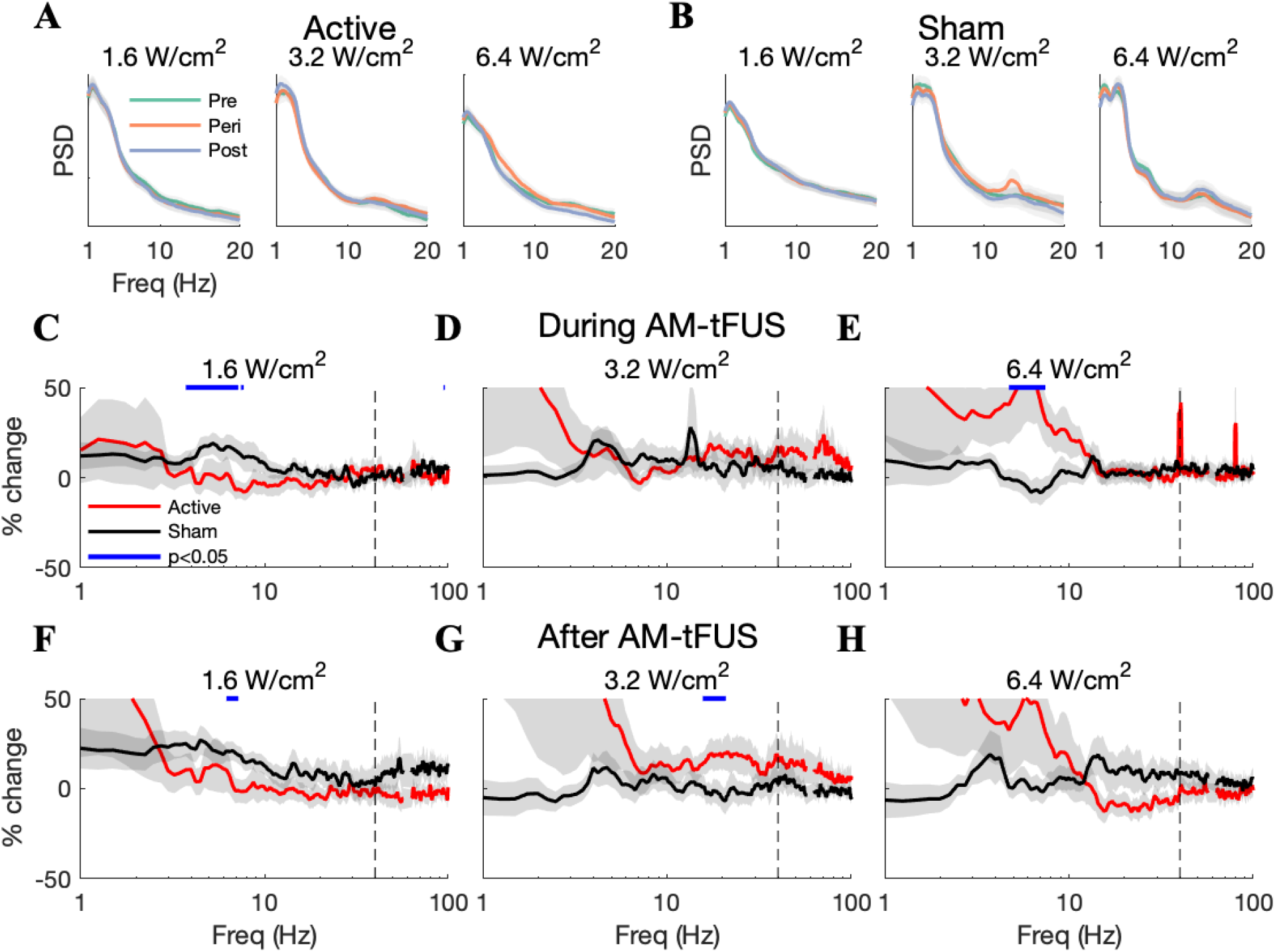
AM-tFUS modulates LFP spectral power in an intensity and frequency-dependent manner in area CA1. Panels replicate Fig 4 of the main text, except that here we show results from the CA1 region of the hippocampus. As in the dentate gyrus, a bidirectional effect on theta power was observed. On the other hand, the significant high-frequency clusters after AM-tFUS were not resolved here. Note that the presence of large increases during high-intensity AM-tFUS at 40 and 80 Hz likely indicates physical displacement of the probe.

**Figure S2:**
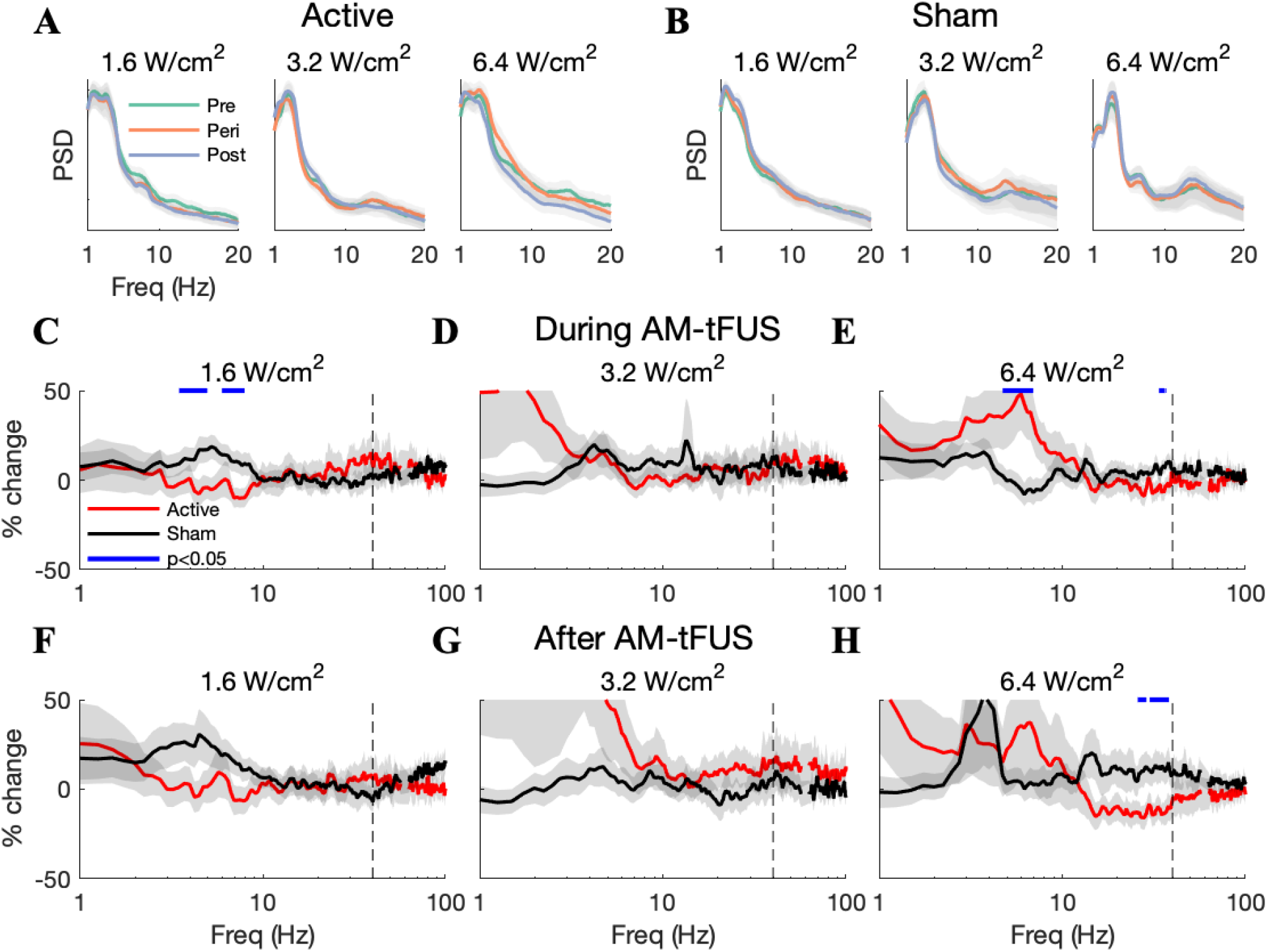
AM-tFUS modulates LFP spectral power in an intensity and frequency-dependent manner in area CA3. Panels replicate Fig 4 of the main text, except that here we show results from area CA3. Unlike in the dentate gyrus and CA1, a significant reduction in the theta band was not resolved after low-intensity AM-tFUS. Note also the significant reductions in gamma power (>30 Hz) during and after high-intensity AM-tFUS.

**Figure S3:**
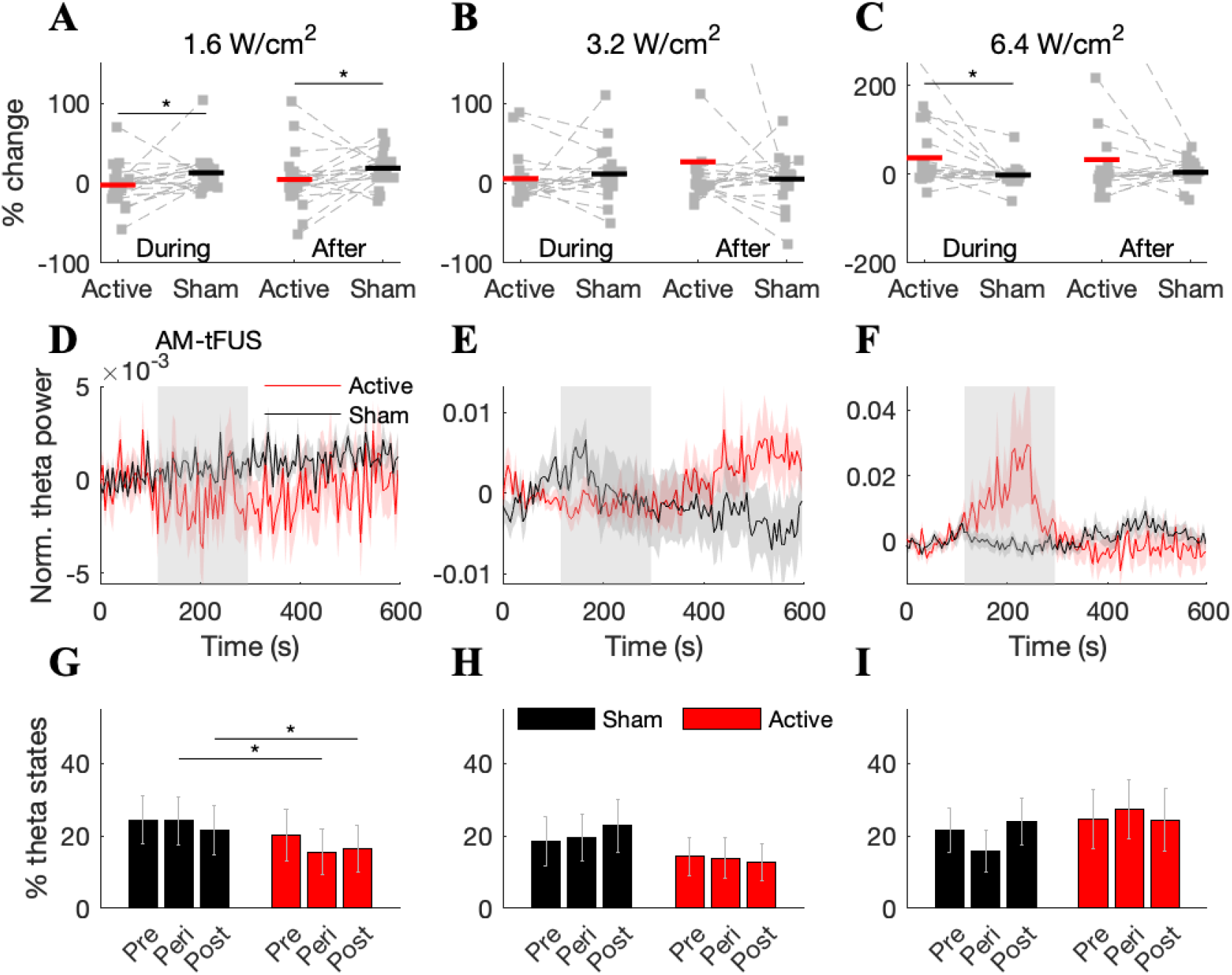
AM-tFUS produces a bidirectional modulation of theta power in area CA1. Panels replicate Fig 5 of the main text but are here shown for CA1. Similar to what was found in the dentate gyrus, the reduction in theta power during low-intensity AM-tFUS outlasted the stimulation. On the other hand, the increased theta power during high-intensity AM-tFUS was not resolved following stimulation.

**Figure S4:**
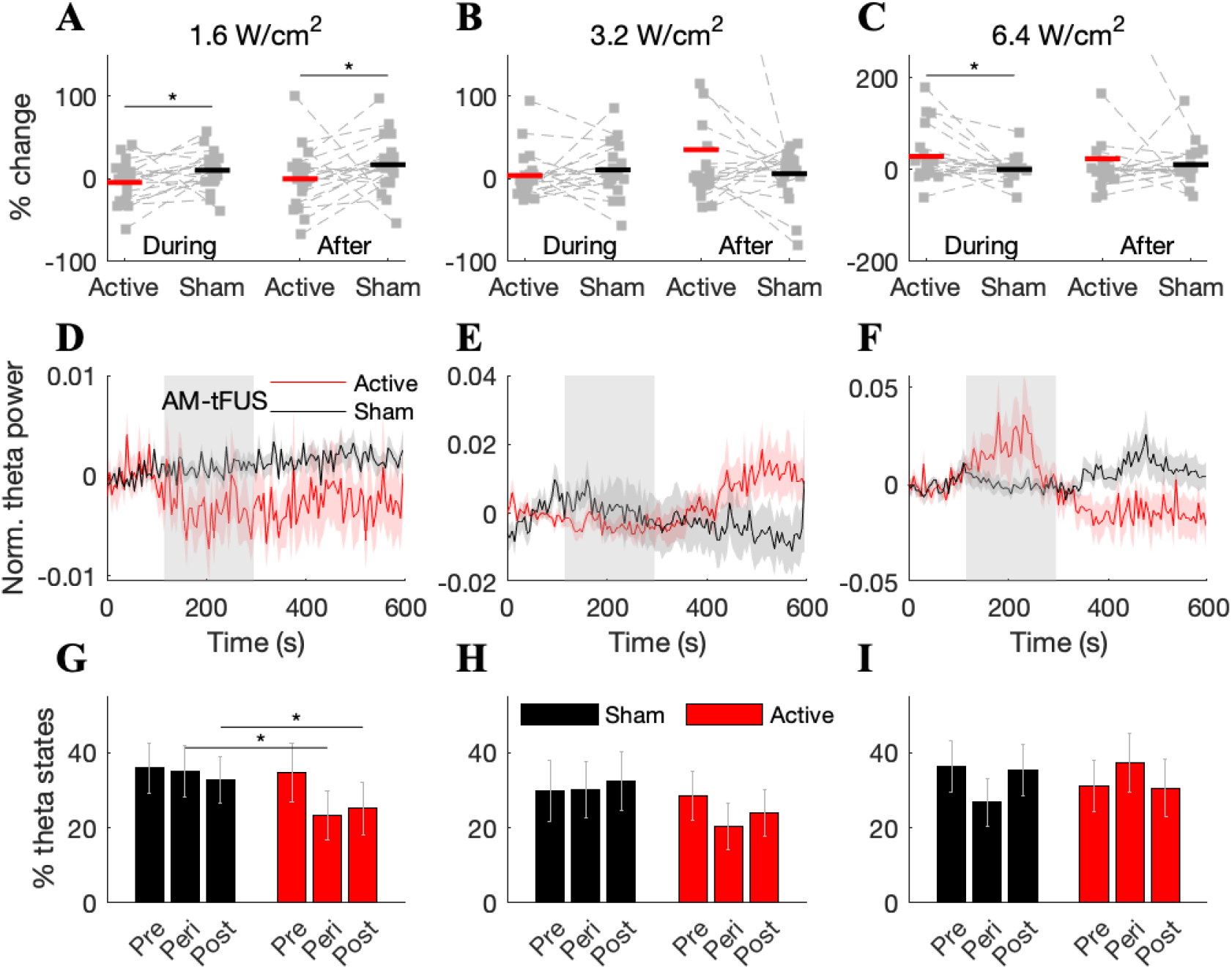
AM-tFUS produces a bidirectional modulation of theta power in area CA3. Panels replicate Fig 5 of the main text but now show data from hippocampal region CA3. The results are qualitatively similar to those found in both the dentate gyrus as well as area CA1.

